# Natural killer cells mediated antibody dependent cellular cytotoxicity response in acute and recovery phases of chikungunya

**DOI:** 10.64898/2026.01.01.697330

**Authors:** Diptee Trimbake, Mohini Ganu, Ashok Ganu, Anuradha S Tripathy

## Abstract

**Background:** Neutralizing antibody is reported to correlate with protection from symptomatic chikungunya disease. In addition to direct virus neutralization, virus-specific antibodies can also protect through antibody effector functions via antibody-dependent cellular cytotoxicity (ADCC) mechanism.

**Methods:** We have assessed NK cell-mediated chikungunya virus (CHIKV) specific ADCC response in 56 acute, 56 prolonged, 44 chronic chikungunya arthritis patients, 23 recovered individuals from chikungunya (CHIK) & 50 healthy controls using a flow cytometry-based antibody-dependent heterologous NK cell activation assay with ADCC surrogate markers CD107a & IFN-γ.

**Results:** Persistent ADCC response in recovered individuals spanning over a duration of 3 months to 18 years post-infection and progressive decline in ADCC response in chronic chikungunya arthritis patients over the period of time were observed. ADCC response correlated with lower plasma viral load in acute patients (*p*=0.022, r=-0.709). Recovered individuals elucidated a positive correlation between ADCC response and plasma anti-CHIKV IgG and IgG3 antibody levels (IgG: r=0.4503, IgG3: r=0.4583, *p*-value<0.01 in each).

**Conclusion:** Overall, the current study establishes the prognostic role of NK cells-mediated ADCC response in chikungunya infection. In a nutshell, these findings provide a base for future antibody-based therapeutics for chikungunya utilizing IgG3 antibody that has functional ADCC capacity.

**Graphical Abstract:** 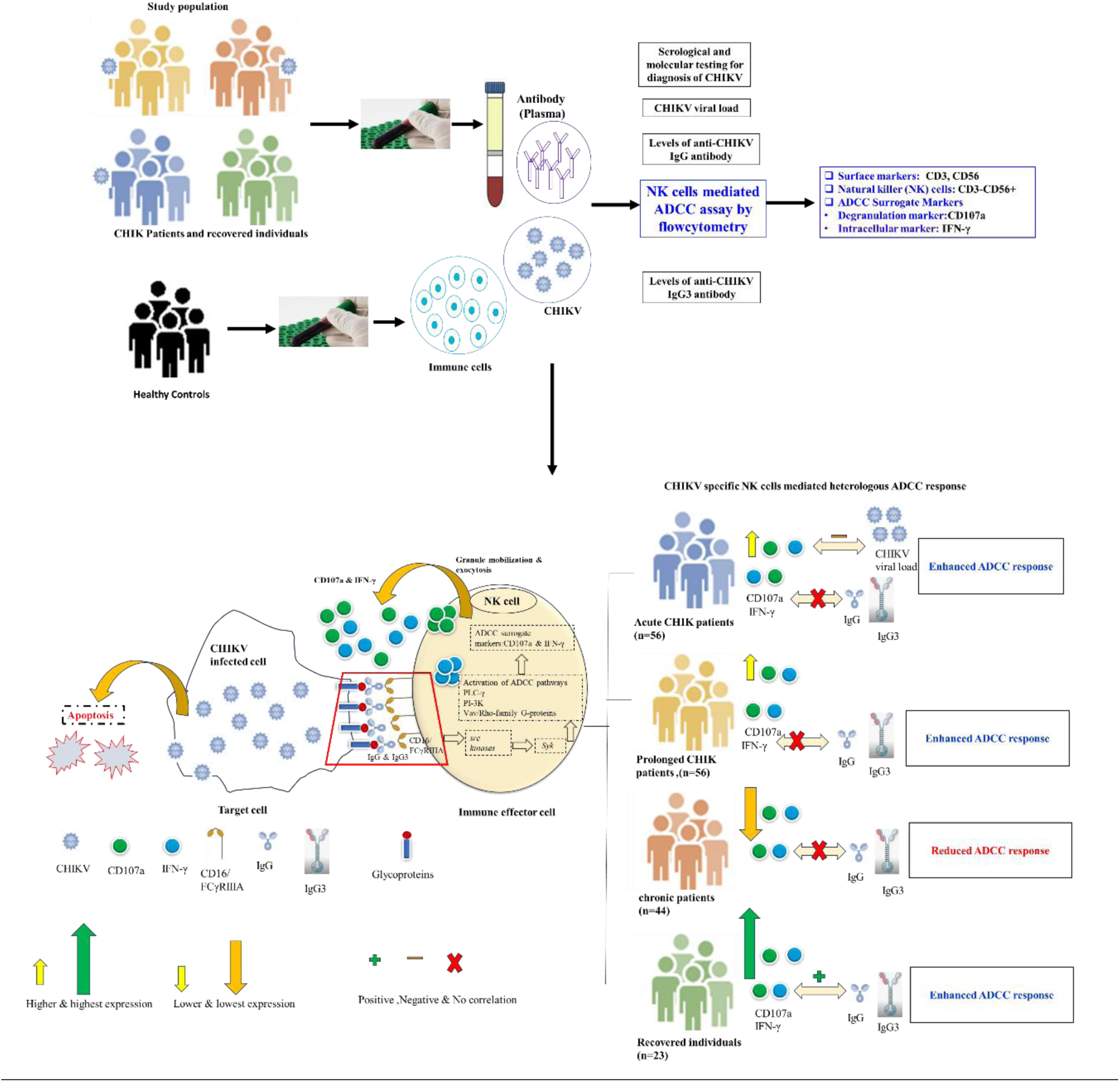

**Highlights:**

- NK cells play key role in chikungunya infection
- NK cells mediated ADCC response remain unexplored in chikungunya
- Anti-CHIKV IgG3 antibody dependent activation of the NK cells observed
- Efficient NK cells mediated ADCC response detected in acute & recovery phases of chikungunya
- IgG3 with functional ADCC capacity can direct Ab-based therapeutics design in chikungunya

## 1. Introduction

Chikungunya (CHIK) is a neglected tropical disease caused by chikungunya virus (CHIKV), designated as a WHO Blueprint priority pathogen [1–2]. Though chikungunya presents as a self-limiting disease, it also leads to chronic manifestation that can affect an estimated 40% of the individuals infected with chikungunya virus [3–4]. It is characterized by high fever (>39°C) during acute illness followed by prolonged polyarthralgia lasting from weeks to years [5]. Chikungunya treatment is mostly supportive and relies on anti-inflammatory drugs and analgesics, which aimed at symptom relief [4–6]. Very recently, two chikungunya vaccines have been authorized for use. VLA1533/Ixchiq, a live-attenuated vaccine, was approved in November 2023 for use in adults aged 18 and above in US, Europe, and Canada. Vimkunya, a virus-like particle vaccine, received approval in February 2025 for use in individuals aged 12 and above in the US. Both are recommended by the FDA and CDC for travellers to outbreak regions and high-risk groups [7].

The processes by which innate and adaptive immune responses mediate the outcome of chikungunya infection are not sufficiently understood. Data over the years suggest that the rapid development of antiviral antibody and T-cell responses are associated with better chikungunya outcome [8–9]. Neutralizing antibodies provide protection by interfering with viral entry, while non-neutralizing antibodies contribute to antiviral defence via Fc-mediated mechanisms such as complement-dependent cytotoxicity, antibody-dependent cellular cytotoxicity (ADCC), and phagocytosis [10]. NK cells play a pivotal role by mediating ADCC through recognition of viral antigens on infected cell surfaces and with the induction of the release of cytotoxic granules resulting in the killing of infected cells [11–12]. NK cells express CD16 (FcγRIIIa), which binds to IgG1 and IgG3.Upon recognition of antibody-coated viral antigens, CD16 triggers signalling pathways leading to perforin/granzyme release and IFN-γ secretion [13]. ADCC is assessed by measuring IFN-γ and/or CD107 expression following antigen-antibody stimulation [14]. ADCC serves as a valuable marker for vaccine and therapeutic efficacy, with elevated ADCC linked to improved clinical outcomes and reduced disease severity [14]. Deceased influenza patients had low ADCC, while vaccination boosted it against H5 and H7, reinforcing its protective role [15]. ADCC aids in early dengue control [16]. ADCC is used as a tool for the evaluation of efficacy of vaccines and antibody-based therapeutics against Ebola virus disease [17]. High ADCC and low Env-specific IgA are linked with reduced HIV risk, highlighting ADCC as a key correlate of protection in the RV144 HIV vaccine [19]. Recovered COVID-19 individuals had higher ADCC than severe and deceased patients [20]. SARS-CoV-2 infection or vaccination induces strong NK cell-mediated ADCC [21].

We have previously reported that acute self-limiting chikungunya patients exhibited higher NK cell percentages, higher expressions of CD107a, and perforin [22]. In contrast, chronic chikungunya arthritis patients showed lower CD107a and perforin expressions indicating impairment of NK cells, thus shedding light on the critical function of NK cells in the pathogenesis and/or protection of chikungunya [23–24]. It seems that the contributions of NK cells to chikungunya disease and protection is evident, though the full spectrum of NK cell actions in chikungunya remains to be elucidated.

ADCC is typically mediated by NK cells, but current knowledge of the mechanisms by which NK cells and antibodies can protect against CHIKV infection and disease or, conversely, contribute to disease development, remain obscure. The challenge remains to identify immune correlates of protection against CHIKV infection or vaccination. This knowledge gap has hampered the rational design of effective therapeutic interventions. Given the importance of NK cells in chikungunya, assessing ADCC involvement in chikungunya is crucial. In the current study, we have evaluated chikungunya virus specific heterologous NK cells mediated ADCC response in acute chikungunya patients (n=56), prolonged chikungunya patients (n=56), chronic chikungunya arthritis patients(n=44), recovered individuals (n=23) from chikungunya and in chikungunya naïve healthy individuals (n=50) from Maharashtra, India.

## 2. Materials and Methods

### 2.1 Ethical approval

The current study was approved by the Institutional Ethics Committee for Research on Humans (Approval No. NIV/IEC/2018/November/D-36) as per the guidelines of the Indian Council of Medical Research, New Delhi. Written informed consents were obtained from all the study participants in accordance with the Declaration of Helsinki.

### 2.2 Study subjects and sample processing

The study population included a total of 156 chikungunya patients (56 acute, 56 prolonged, and 44 chronic chikungunya arthritis patients), 23 recovered individuals and 50 chikungunya naïve healthy individuals. The recovered individuals were enrolled during the follow-up camps organized at Sanjeevan Hospital, Latur, Maharashtra. Control samples were collected from the blood-donation camps. Whole blood was used to isolate peripheral blood mononuclear cells (PBMCs), while plasma was utilized for serological, molecular, and NK cell-mediated ADCC studies. The diagnosis of CHIKV has been confirmed through the detection of IgM antibodies in ELISA and/or RT-PCR analysis.

Patients were categorized based on the symptoms and number of day’s post-onset of illness as follows:**1. Acute chikungunya patients** had acute febrile polyarthritis (≤ 15 days duration) and were positive for anti-CHIKV IgM antibodies and/or CHIKV RNA, and positive/negative for anti-CHIKV IgG antibodies. **2. Prolonged chikungunya patients** were positive/negative for anti-CHIKV IgM/IgG antibodies and presented polyarthritis/arthralgia that persisted between the duration of 15 days and 3 months.**3. Chronic chikungunya arthritis patients** were positive/negative for anti-CHIKV IgM antibodies, were positive for anti-CHIKV IgG antibody and presented with inflammatory symmetric polyarthritis/arthralgia that persisted for more than 3 months following an episode of chikungunya.**4. Recovered** i**ndividuals** were negative for anti-CHIKV IgM and positive for anti-CHIKV IgG antibody and remained asymptomatic for at least three months following the acute phase of chikungunya infection. **5. Chikungunya naïve healthy individuals** reported absence of any previous history of chikungunya and negative for anti-CHIKV IgM and IgG antibodies. Exclusion criteria for the study included individuals undergoing steroid treatment, rheumatoid arthritis patients positive for RA factor, anti-cyclic citrullinated peptide (CCP) antibody, anti-nuclear antibody (ANA) with elevated serum uric acid levels [24]. The study design is illustrated in **Supplementary Figure 1.** The number of samples from each category used for different assays is displayed in **Supplementary Figure 2.** Detailed characteristics are provided in **Table 1**.

**Table 1.** Characteristics of the study groups.

| **Parameters** | **Acute**<br>**chikungunya**<br>**patients** | **Prolonged**<br>**chikungunya**<br>**patients** | **Chronic chikungunya**<br>**arthritis patients** | **Recovered**<br>**individuals** | **Healthy**<br>**controls** |
| --- | --- | --- | --- | --- | --- |
| **Study population** | n=56 | n=56 | n=44 | n=23 | n =50 |
| **Sex ratio**<br>**(male: female)** | 0.21(10/46) | 0.27(12/44) | 0.37(12/32) | 1.5(14/9) | 4(40/10) |
| **Age (years)** | 47(23-72) | 44(18-73) | 42(19-68) | 48(23-66) | 24(19-55) |
| **POD** | 6 days<br>(1day-15 days) | 38 days<br>(16 days-90 days) | 3 months<br>(3 months-17 years) | 1 year 2 months#<br>(3 months -18 years) | NA |
| **Anti-CHIKV IgM** | Positive, n= 46 | Positive, n= 56 | Positive, n= 20 | NA | All samples<br>were negative |
|  | Negative, n=6 |  | Negative, n=24 |  |  |
|  | Indeterminate, n=4 |  |  |  |  |
| **CHIKV nested PCR** | Positive, n=10 | ND | ND | ND | NA |
| **Viral load (copies/ml)** | 4880<br>(263200-999200000) | ND | ND | ND | NA |
| **Anti-CHIKV IgG** | Positive, n=46 | Positive, n= 53 | Positive, n = 44 | Positive, n=23 | All samples<br>were negative |
|  | Negative, n= 10 | Negative, n=3 |  |  |  |
| **Anti-CHIKV IgG titer** | 1600(100-6400) | 3200(200-12800) | 1600(200-12800) | 6400(200-102400) | NA |
| **Anti-CHIKV IgG3**<br>**titer** | 400(100-6400) | 800(100-6400) | 800(100-3200) | 12800(800-51200) |  |

### 2.3 Serological and molecular testing for diagnosis of CHIKV infection

All the samples were screened by an in-house ELISA for antibodies against CHIKV and DENV (anti-CHIKV/DENV IgM and IgG antibodies) as previously described [22–23].

### 2.4 Viral load quantification

CHIKV viral load (copies/ml) was determined from the plasma samples of acute patients using extracted RNA, primers and probes based on CHIKV E3, as described earlier [22,26–27].

### 2.5 Purification and inactivation of CHIKV by β-propiolactone (BPL)

CHIKV isolation, purification and inactivation with BPL (Sigma, USA) was carried out following the previously described protocol [22]. Subsequent to protein estimation, the whole, inactivated chikungunya virus was used as recall antigen in the heterologous NK cell-mediated ADCC experiments.

### 2.6 NK cells mediated heterologous ADCC assay

Anti-CHIKV-specific heterologous ADCC activity in the plasma samples of the study participants was determined using a NK cell activation assay [14,28–29]. Whole blood from chikungunya naïve controls were used as the source of NK cells. CD107a and/or IFN-γ expression by activated NK cells were used as surrogate markers for CHIKV-specific stimulation of NK cells in presence of heterologous antibodies. The recent studies demonstrated that ADCC activity observed in NK cell activation assay correlates significantly with functional killing assays such as the rapid fluorometric ADCC assay and other assays [14].

Briefly, 150 µl whole blood from anti-CHIKV IgG negative healthy donor (as a source of NK cells) and 50 µl of heat-inactivated heparin-anticoagulated plasma [final concentration 1:4] from the study participants was stimulated with CHIKV antigen (10μg/ml) in the presence of brefeldin A (10μg/ml, Sigma Aldrich, St. Louis, MO), monensin (0.68μl/ml, BD Biosciences, USA), anti-CD107a APCH7 (clone H4A3, BD Biosciences, USA) and incubated for 6 hrs at 37^0^C in a 5% CO_2_ incubator. Stimulation with purified anti-CD16 antibody (clone-3G8, 5μg/ml, BD Biosciences, USA) served as a positive control. Plasma samples of all study groups incubated with anti-CHIKV IgG negative whole blood in the absence of antigen was considered as a negative control for the respective sample. After incubation, staining was carried out for CD3−CD56dim NK cells using fluorescent tagged antibodies, CD3 PerCp (clone SK7, BD Biosciences, USA) and CD56 PECy7 (clone NCAM16.2, BD Biosciences, USA) antibodies. RBCs were lysed using 1X BDFACS lyse solution (BD Biosciences, USA) and cells were permeabilized using 1X BD PERM-II solution (BD Biosciences, USA). Cells were washed and processed for intracellular staining using anti-IFN-γ APC [Clone 25723.11 (RUO (GMP), BD Biosciences, USA] antibody, followed by washing with 1X PBS and fixing with 1% paraformaldehyde. The cells were then acquired on FACS ARIA-II (BD Biosciences, USA). Single colour compensation was performed prior to acquisition of samples in the FACS Aria II flow cytometer. For each experiment, 50,000 events were recorded. The lymphocytes were gated by forward and side scatter, identifying NK cells as CD3-CD56+. Gating strategy for antibody-dependent NK cell activation is described in **Supplementary Figure 3**. The data were analysed using FACS Diva software version 8(BD Biosciences, USA).

The ADCC activity was represented as the percentage of any activated NK cells expressing only CD107a/ both CD107a and IFN-γ/ IFN-γ, **Figures 1**. The percent NK cell activation seen in the unstimulated control of each sample was subtracted from the CHIKV stimulated to measure CHIKV induced NK cell activation. The response was considered to be positive if the percentage of activated NK cells expressing both CD107a and IFN-γ of CHIKV stimulated was three times greater than the percent activated NK cells of the respective unstimulated control.

**Figure 1.**
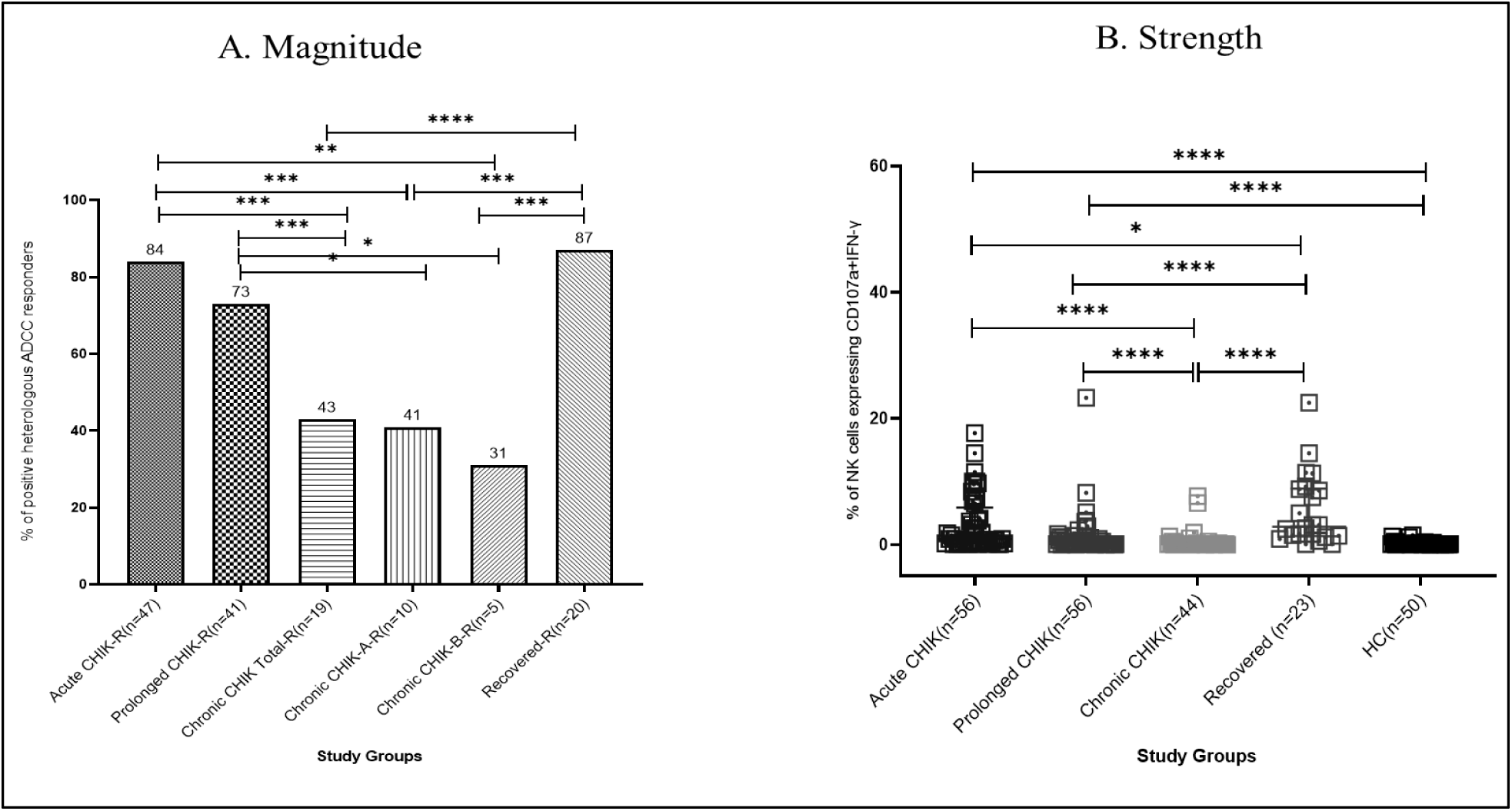
NK cells mediated heterologous ADCC response in chikungunya patients and recovered individuals. *CHIK: Chikungunya, R: responders, A: POD<1 year, B: POD>1 year, Bar graphs representing % of positive heterologous ADCC responders. ADCC positive responders: Test >3*NC, ,2*2 chi-square (Fisher’s exact) test, *p-value <0.05, **p-value <0.005, ***p-value <0.0001, ns=non-significant*

### 2.7 Estimation of plasma anti-CHIKV IgG and IgG3 antibody levels by ELISA

Monoclonal antibodies targeting mouse anti-human IgG antibody subclasses (anti-human IgG3) were employed to detect CHIKV-specific IgG and IgG3 antibodies using an ELISA-based assay [30]. Each assay included three negative and two positive controls. The ELISA cut-off was set at three times the mean absorbance of negative controls. IgG3 titres were defined as the highest plasma dilution with OD ≥ cut-off. A log10 titre of 0.301 was used for negative plasma in the analysis.

### 2.8 Software and statistical analyses

The statistical analyses were performed on IBM SPSS Statistics 25 software (SPSS Inc., Chicago, IL, USA) and GraphPad Prism 8 software (GraphPad, San Diego, CA, USA). All the data are expressed as the median (IQR). *Mann-Whitney* test was used for group comparisons. Fisher’s exact test was used for the binary data (ADCC responders and non-responders, age below and above 50 years, anti-CHIKV IgG antibody positives and negatives). One-sided, two-sample test of proportions was applied to compare gender-wise differences in the ADCC responses (percent male and female responders). Spearman rank correlation test was used to assess the correlation between ADCC responses of the NK cells expressing surrogate markers and i) viral load, ii) total anti-CHIKV IgG and IgG3 antibody levels. *P* values <0.05 were considered significant. The dot plots were generated on GraphPad Prism 8 software (GraphPad, San Diego, CA, USA).

## 3. Results

### 3.1 Characteristics of the study population

The study population comprised of five distinct groups with a total of 229 participants. **Table 1** presents the detailed characteristics of the study groups.

### 3.2 NK cells mediated heterologous ADCC response

CHIKV-specific ADCC response was evaluated using an NK cell activation assay, with CD107a and IFN-γ expressions serving as surrogate markers.

#### 3.2.1 Magnitude of NK cells mediated heterologous ADCC response

Forty seven of 56 (84%) acute chikungunya patients, 41of 56 (73%) prolonged chikungunya patients, 19 of 44 (43%) chronic chikungunya arthritis patients and 20 of 23 (87%) recovered individuals exhibited antibody-dependent heterologous NK cells response against CHIKV antigen. Ten of 24 (41%) chronic chikungunya arthritis patients with POD < 1 year and 5 of 16 (31%) chronic chikungunya arthritis patients with POD >1 year demonstrated antibody-dependent heterologous NK cells response against CHIKV antigen (*p* value<0.001 in each) **Figure 1A**.

#### 3.2.2 Strength of NK cells mediated heterologous ADCC response

*Mann Whitney U* test was used to determine the strength of ADCC response. Acute and prolonged chikungunya patients displayed significantly higher CHIKV specific NK cell mediated heterologous ADCC response compared to chronic chikungunya arthritis patients and healthy controls. Acute, prolonged chikungunya patients and healthy controls presented significantly higher CHIKV specific NK cell mediated heterologous ADCC response compared to chronic chikungunya arthritis patients. Recovered individuals displayed significantly higher CHIKV specific NK cell mediated heterologous ADCC response compared to acute, prolonged and chronic chikungunya arthritis patients [**Dual expressions of CHIKV specific CD107a and IFN-γ on NK cells:** acute chikungunya patients: 1.7(0.1-33.5), prolonged chikungunya patients: 0.3(0.0-23.3), chronic chikungunya arthritis patients: 0.1(0.0-7.7), recovered individuals: 2.8(1.3-8.9), and healthy controls: 0.1(0.0-1.5), *p*=0.0001 in each] **Figure 1B**.

It is noteworthy that acute chikungunya patients and recovered individuals demonstrated significantly higher heterologous antibody dependent activation of NK cells with expression of single ADCC surrogate marker CD107a or IFN-γ compared to the chronic chikungunya arthritis patients. [**Expressions of CHIKV specific NK cells with CD107a:**acute chikungunya patients: 4.3(0.1-32.0), chronic chikungunya arthritis patients: 0.1(0.0-16.6), recovered individuals: 0.9(0.1-13.4) and healthy controls: 0.1(0.0-6.4), **Expression of CHIKV specific NK cells with IFN-γ**: acute chikungunya patients:1.3(0.1-13.7), chronic chikungunya arthritis patients: 0.1(0.0-7),recovered individuals:1.8(0.7-5.6) and healthy controls: 0.1(0.1-0.2) *p*=0.0001 in each] **Figure 2**.

**Figure 2.**
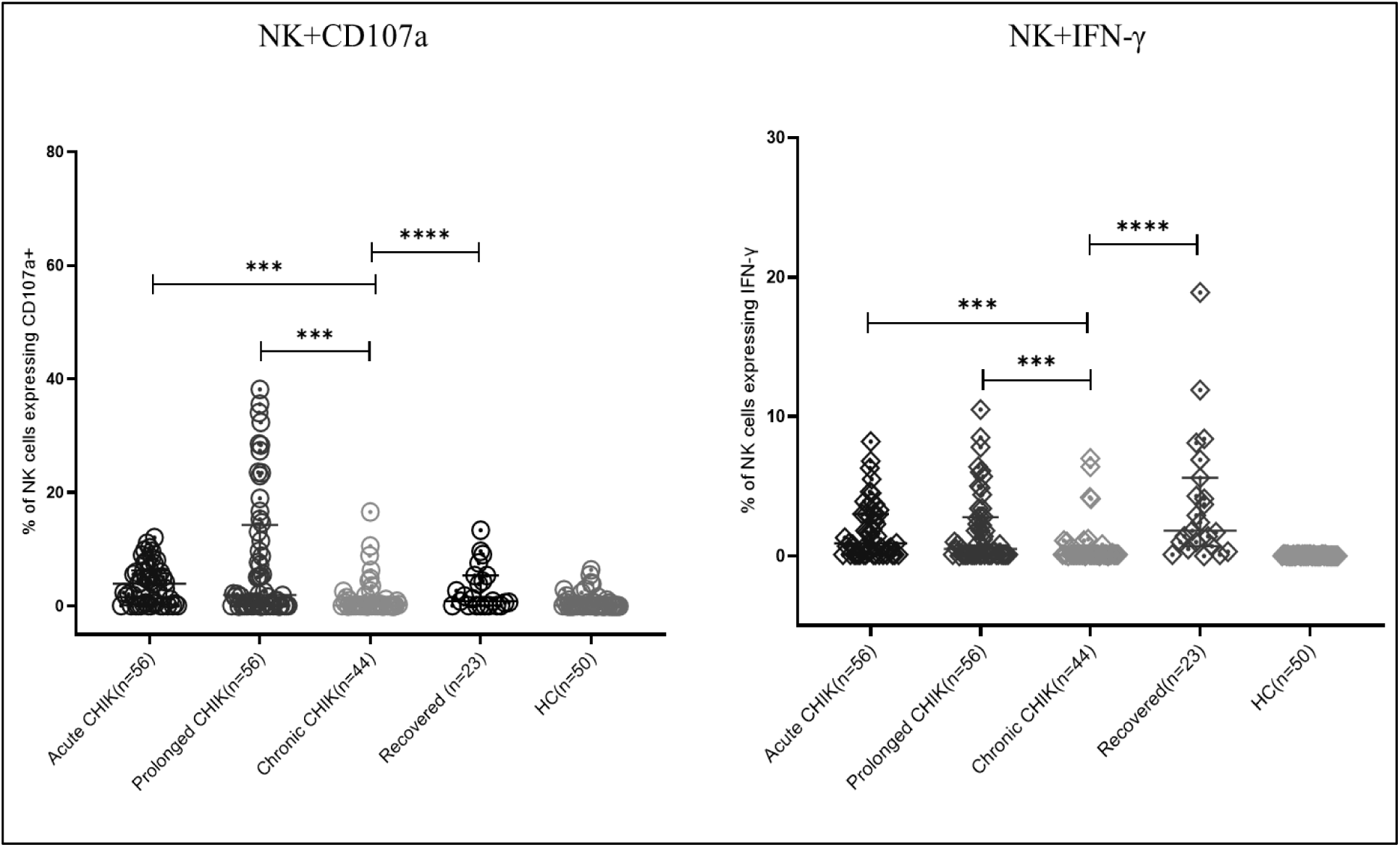
ADCC response with expressions of single ADCC surrogate marker. *The dots represent individual values and data represented as median (IQR) values, Mann Whitney U test, *p-value <0.05, **p-value <0.005, ***p-value <0.0001, ns=non-significant*

The data demonstrate stronger recruitment of activated NK cells in acute chikungunya patients and in recovered individuals via heterologous ADCC response.

### 3.3 Association of heterologous ADCC response with age

Based on the age, the study participants were categorized into two groups, less than 50 and more than 50 years. In the below 50 years of age category, recovered individuals showed significantly higher CHIKV specific ADCC response compared to acute, prolonged and chronic chikungunya arthritis patients [**Expressions of CHIKV specific CD107a and IFN-γ on NK cells:** acute chikungunya patients:1.7(0.1-33.5), prolonged chikungunya patients:0.1(0.0-8.2), chronic chikungunya arthritis patients :0.1(0.0-7.7), recovered individuals:3.1(1.3-9.2), *p* value<0.05 in each).

### 3.4 Association of heterologous ADCC response with gender

Chi-square (Fisher’s exact) one-sided two-sample test of proportions demonstrated that ADCC response in acute, prolonged chikungunya, chronic chikungunya arthritis patients and recovered individuals was independent of gender.

### 3.5 Levels of anti-CHIKV IgG and IgG3 antibody

Recovered individuals had significantly higher levels of total anti-CHIKV IgG and IgG3 antibodies compared to acute and chronic chikungunya arthritis patients (*p* value<0.0001 in each), **Figure 3**.

**Figure 3.**
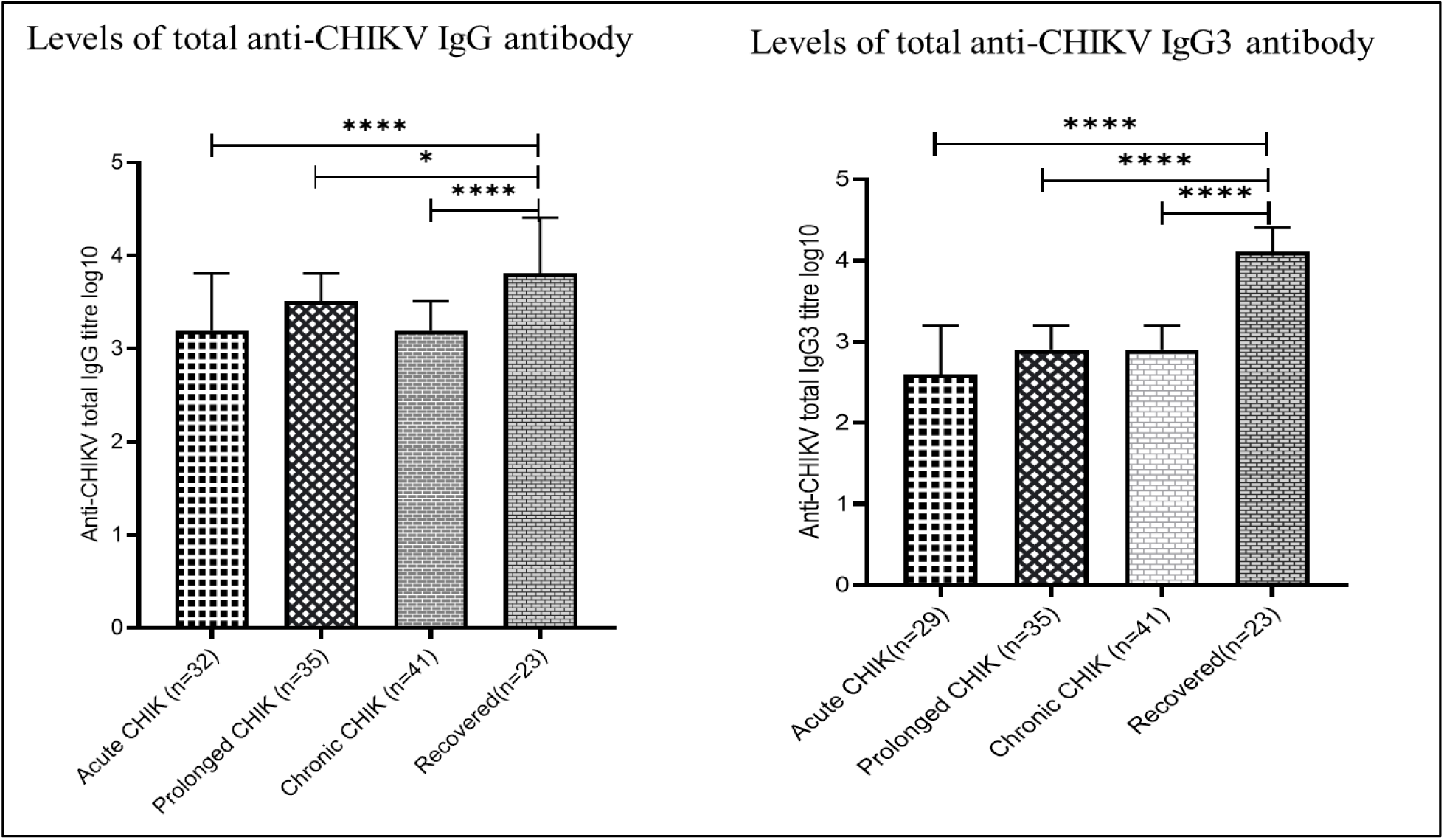
Levels of anti-CHIKV IgG and IgG3 antibody. *CHIK: chikungunya, Bar graphs representing levels of anti-CHIKV IgG and IgG3 antibody. Data shown as median (IQR) values. Mann Whitney U test*p-value <0.05, **p-value <0.005, ***p-value <0.0001, ns=non-significant*

### 3.6 Association of heterologous ADCC response with anti-CHIKV IgG antibody status

Among the anti-CHIKV IgG positives, 38 of 46 (82.6%) acute chikungunya patients, 39 of 53

(73.5%) prolonged chikungunya patients, 19 of 44 (43.2%) chronic chikungunya arthritis patients, and 20 of 23 (87%) recovered individuals demonstrated ADCC response against CHIKV antigen. Among the anti-CHIKV IgG negatives, 9 of 10 (90%) acute chikungunya patients, 2 of 3 (66%) prolonged chikungunya patients demonstrated ADCC response against CHIKV antigen, **Supplementary Figure 4.**

ADCC responders had significantly higher levels of total anti-CHIKV IgG and IgG3 antibodies compared to non-responders in the acute chikungunya patients’ group (*p* value<0.01) **Figure 4**.

**Figure 4.**
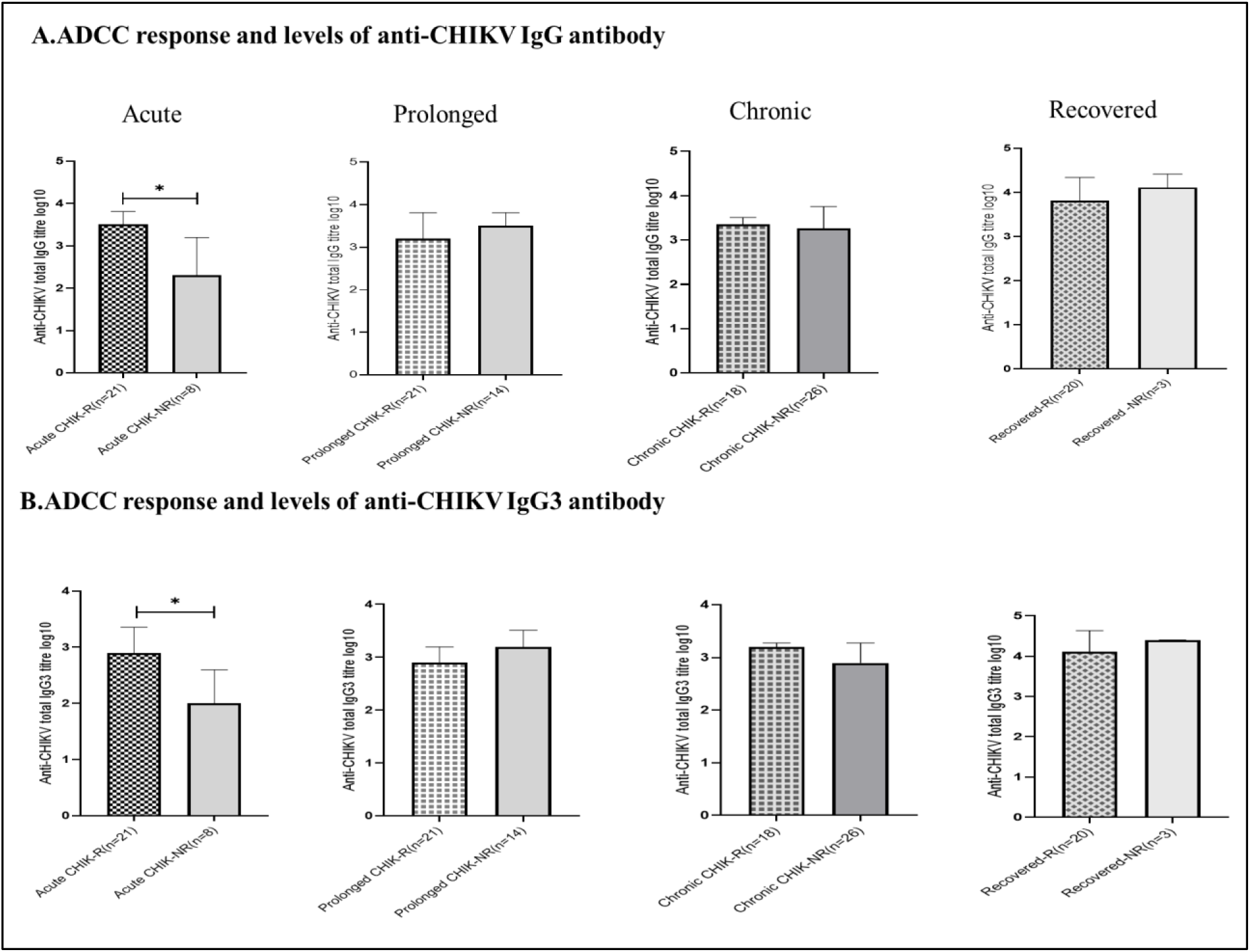
ADCC response and the levels of anti-CHIKV IgG & IgG3 antibodies. *CHIK: Chikungunya, R: ADCC responders, NR: ADCC positive responders: Test >3*NC, Bar graphs representing levels anti-CHIKV IgG and IgG3 antibody. Data shown as median (IQR) values. Mann Whitney U test*p-value <0.05, **p-value <0.005, ***p-value <0.0001, ns=non-significant*

### 3.7 Correlation analysis

#### 3.7.1 ADCC response and CHIKV viral load

A negative correlation was detected between the level of ADCC response and plasma CHIK viral load in acute chikungunya patients (r=-0.709, *p*=0.022) **Figure 5**.

**Figure 5.**
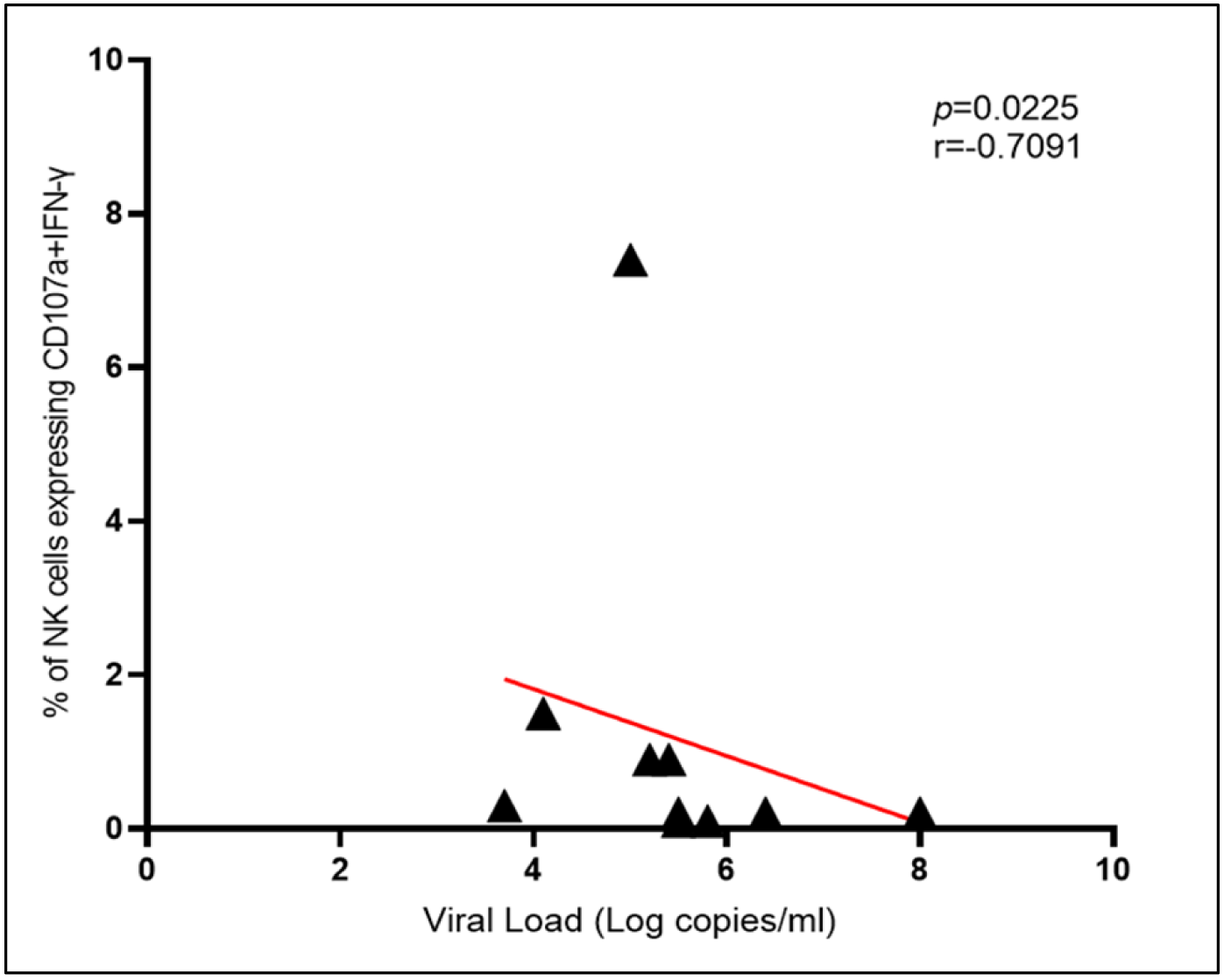
Correlation between heterologous ADCC responses and CHIK viral loads. *The scatter plots show the Spearman rank correlation to determine the NK cells mediated heterologous ADCC response with ADCC markers CD107a and IFN-γ (Y-axis) with the plasma CHIK viral load (log copies/mL) (X-axis) in acute chikungunya patients, Spearman correlation coefficient (r) = -0.5363, p = 0.0225. Dashed line represents 95% confidence intervals around the correlation*

#### 3.7.2 ADCC response and anti-CHIKV IgG & IgG3 antibody levels

Recovered individuals showed a positive correlation between heterologous CHIKV specific ADCC activation of NK cells (CD107a and IFN-γ) and plasma anti-CHIKV IgG and IgG3 antibody levels (recovered individuals: IgG: r=0.4503, IgG3: r=0.4583, *p* value<0.01) **Figure 6**.

**Figure 6.**
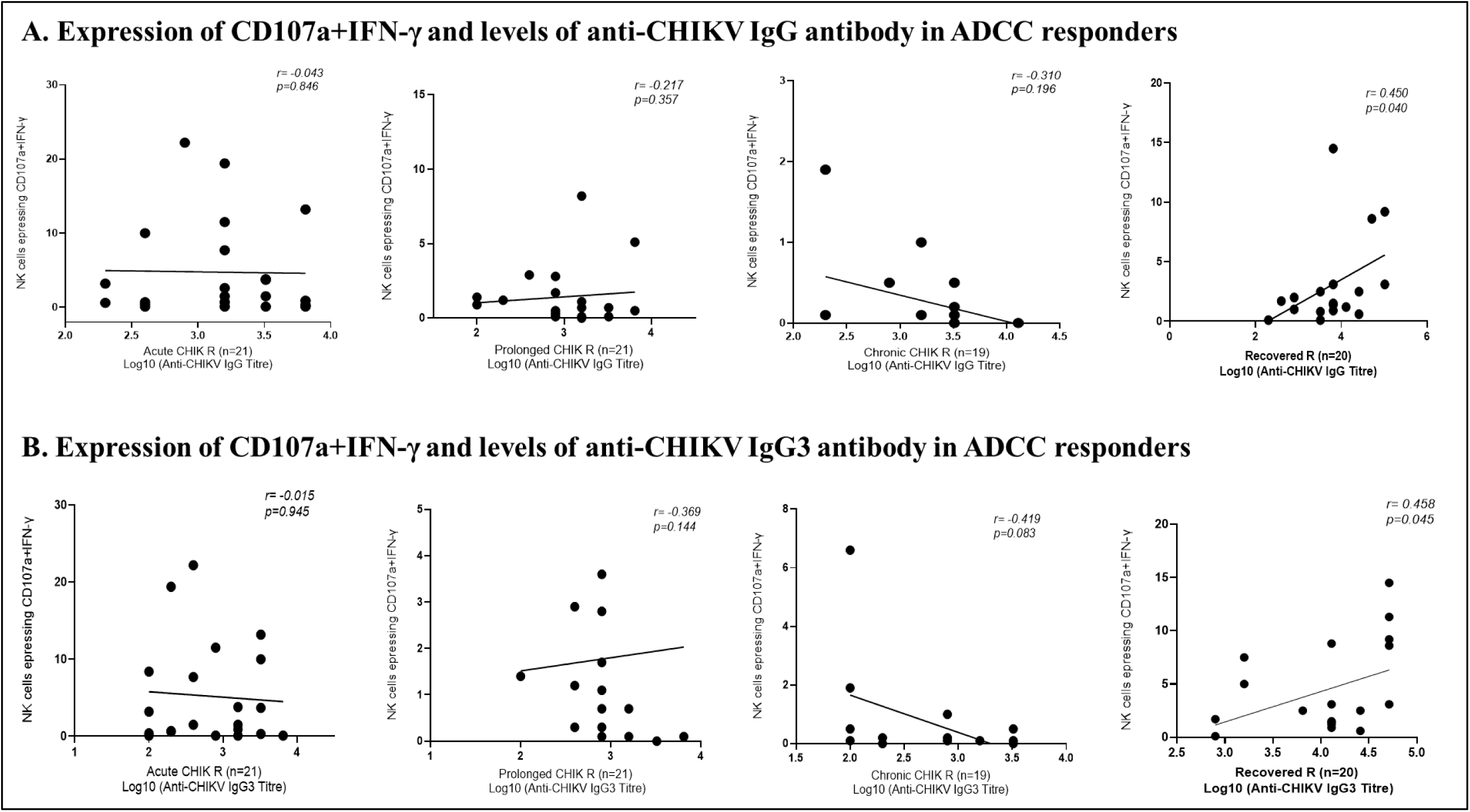
Correlation between ADCC responses and total anti-CHIKV IgG and IgG3 antibody levels. *The scatter plots show the Spearman rank correlation to determine the NK cells mediated heterologous ADCC response with ADCC markers CD107a+IFN-γ (Y-axis) in chikungunya patients and recovered individuals with total anti-CHIKV IgG and IgG3 levels (log 10) (X-axis), Spearman correlation coefficient (r). A p-value <0.05 is considered significant. Dashed line represents 95% confidence intervals around the correlation*

## 4. Discussion

The CHIKV vaccines, IXCHIQ (formerly VLA1553) and VIMKUNYA (formerly PXVX0317), were licensed in the US and the European Union between 2023 and 2025 through accelerated pathways [7]. Detection of IgG antibodies along with the neutralizing antibodies are considered the presumptive sero-protective response [31]. Fc-mediated functions of antibodies, such as ADCC have been correlated with protection against HIV infection in an RV 144 phase III HIV vaccine trial [19]. In SARS-CoV-2, elevated ADCC responses have been reported in recovered individuals compared to those with severe disease or fatal outcomes with persistence documented up to 12 months following infection or vaccination [20]. MV-CHIK, a measles-vectored, live-attenuated recombinant chikungunya vaccine elicited robust antibody effector functions, prominently featuring ADCC activity characterized by CD107a-mediated degranulation and IFN-γ cytokine release, indicative of potent Fc-dependent antiviral immunity [32–33]. Our previous studies have highlighted the prognostic role of NK cells in modulating CHIKV pathogenesis and recovery [22–25]. However, the role of Fc-mediated effector functions, particularly NK cells-driven ADCC, remains insufficiently characterized [34]. The current study provides a comprehensive evaluation of NK cell-mediated ADCC responses across different stages of chikungunya that may help to identify the role of ADCC and clarify the contribution of Fc-mediated effector functions to support the development of advanced immunotherapeutic strategies leading to a better clinical outcome.

Thanapati et al. reported higher percentages of peripheral NK cells, higher MFI of CD107a+NK cells in the acute patients than in controls, suggesting that the degranulation capacity per NK cell in the acute CHIKV patients is higher, and enhanced percentage of perforin/ IFN-γ + NK cells in the acute phase indicated the regulatory role of NK cells in acute CHIKV infection **[22].** Our current data of higher antibody dependent heterologous NK cells activation with ADCC surrogate markers CD107a and IFN-γ in the acute (84%) and prolonged chikungunya patients (73%) compared to healthy controls (0%), thus indicating that the recruitment of NK cells through antibody-mediated means may serve for the early control of CHIKV infection. This could be suggestive of early cytolytic ADCC responses reflecting timely FcγR engagement with efficient clearance of infected cells prior to immune dysregulation.

Our earlier observations revealed a positive correlation between NK cells percentage and viral load in the acute-chikungunya patients **[22].** Extending these findings, the present study reports an inverse association between CHIKV viral load and heterologous ADCC responses in the acutely infected chikungunya patients suggesting that CHIKV-specific antibodies may facilitate the regulation of viremia via FcγR-mediated activation of NK cells, as noted in other viral settings **[35].** Previous reports have demonstrated a significant role of ADCC antibodies in the early suppression of secondary DENV-3 viremia **[16]** and in the potential reduction of the latent HIV reservoir **[17].** Collectively, these findings underscore the importance of Fc-mediated NK cell effector functions during the acute phase of CHIKV infection and highlights ADCC as a potential immunological marker for protective antiviral responses that can mediate clearance of infected cells.

Our earlier study has demonstrated impaired NK cell cytotoxicity in chronic chikungunya patients, with marked reductions in perforin and CD107a expressions **[23].** In concordance, the present investigation identified a progressive decline in NK cell-mediated ADCC response in chronic chikungunya patients (43%) over the period of time. This is suggestive of altered NK cell function with decreased cytolytic activity linked to the chronic phase of the disease. The possible cause of this observation could be the suppressed activating receptors (NKp80, NKp46, NKG2D) that may contribute to ADCC impairment, as observed in persistent viral infections including Kaposi’s sarcoma-associated herpesvirus [36], human immunodeficiency virus [37] and chronic HCV carriers [38], highlighting a possible conserved mechanism of immune evasion.

Thanapati et al. observed increased IFN-γ expression on NK cells of recovered individuals from chikungunya compared to those with chronic disease manifestation **[23].** Extending these findings, our current study detected persistent NK cell-mediated ADCC responses in 87% of recovered chikungunya individuals, spanning over a duration of 3 months to 18 years post-infection. These long-lived functional responses suggest a durable immunological imprint, potentially contributing to protective immunity. Longitudinal follow-up of recovered individuals assessing the susceptibility to re-infection and the persistence of ADCC functionality may further clarify its prognostic relevance.

In CHIKV infection, IgG1 and IgG3 play key roles as potent activators of innate immune effector cells in the adaptive immune response, particularly in Fc-mediated effector functions [44]. The binding capacity of IgG1 and IgG3 in ADCC is a critical factor in their effectiveness in eliminating virus-infected cells [13]. However, IgG3 generally has higher affinity than IgG1, due to better accessibility provided by its longer hinge region, which enhances its interaction with FcγRs [42]. The distinct effector function profile of IgG3 positions is a key mediator of protective immunity, particularly in antiviral responses requiring robust Fc-dependent mechanisms [39-40-41]. Its superior affinity for FcγRIIIa/CD16a, early emergence post-infection, superior protection against intracellular pathogens and enhanced ability to activate NK cells underscores its relevance in driving ADCC **[41–42].** Evidence from the RV144 HIV vaccine trial further underscores IgG3’s immunological significance, where elevated IgG3 binding to HIV-1 Env correlated with reduced infection risk and heightened ADCC activity **[19].** Early IgG3 responses aid viral clearance in acute hepatitis C infection, highlighted their protective role **[43].** An early IgG3-dominant response has been linked with high viremia, severe acute symptoms, and more effective viral clearance, leading to reduced risk of chronic arthralgia in chikungunya infection **[44].** However, data from India present a contrasting narrative, with some studies reporting negligible or absent IgG3 levels during acute chikungunya infection **[45].** Conversely, pooled serological analysis of 25 confirmed CHIKV cases revealed IgG3 as the predominant subclass, specifically targeting envelope protein epitopes **[46].** Our study reports detection of lower levels of anti-CHIKV IgG3 antibodies in the chronic chikungunya patients compared to the recovered individuals attributing to a lower affinity of IgG3 for FcγRIIIa/CD16, resulting in the lack of NK cell mediated ADCC response and possible chronicity. MV-CHIK vaccination induces a dominant IgG3 antibody response which persisted even at 6 months as shown by detailed serological analysis of Fab- and Fc-mediated immunity in phase II trial participants **[33].** In a similar line, the present study observed a positive correlation between ADCC response and plasma anti-CHIKV IgG3 levels in the recovered individuals that is suggestive of the role of Fc dependent effector function of IgG3 towards recovery as reported elsewhere **[47].** It is known that the extended hinge and disulfide bonds of IgG3 enhance flexibility and antigen accessibility, boosting Fc receptor engagement and effector functions. This structural adaptability makes IgG3 a promising candidate for antibody-based therapeutics [42]. Taken together, these data advocate for deeper exploration of IgG3 as a candidate for designing antibody therapeutics that harness potent immune effector functions to combat persistent viral infections.

Higher ADCC response in the age group below 50 years in the recovered individuals from chikungunya infection is an important observation. We report that NK cell mediated ADCC response was gender-neutral, thus indicating that ADCC response primarily relies on factors such as antibodies and the recognition of target cells.

Our study elucidated a robust anti-CHIKV IgG3 antibody dependent activation of the NK cells through a well-coordinated and efficient ADCC response in the acute and recovery phases of chikungunya. This study provides a base for future antibody-based therapeutics for chikungunya utilizing IgG3 having functional ADCC capacity. Assessment of the NK cells-mediated ADCC response in the recovered individuals over a period of time may confirm its prognostic role.

## Acknowledgments

We thank Director, ICMR-National Institute of Virology, for all the encouragement. We greatly appreciate all individuals who participated in this study. We sincerely thank the dedicated staff of the blood donation camp for their invaluable support. Special thanks are due to all members of Hepatitis, Chikungunya, and Dengue group for technical help. Ms. Diptee Trimbake, a Ph.D. scholar at ICMR-National Institute of Virology affiliated with Savitribai Phule Pune University, would like to thank the Council of Scientific and Industrial Research (CSIR), New Delhi, India, for providing the Junior and Senior Research Fellowships.

## Author Contributions

The study was conceived and designed by AST, MG, and AG were responsible for the enrolment of study participants, treatment, clinical data and follow up. DT carried out the experiments, performed the analyses and wrote the first draft. The study was carried out with the internal funds from ICMR-NIV, Pune. All co-authors reviewed, edited and approved the final version of the manuscript.

## Data Availability Statement

The data underlying this article are available in the article and in its online supplementary material.

## Ethical approval

The current study was approved by the Institutional Ethics Committee for Research on Humans (Approval No. NIV/IEC/2018/November/D-36) as per the guidelines of the Indian Council of Medical Research, New Delhi.

## Patient Consent Statement

Written informed consents were obtained from all the study participants in accordance with the Declaration of Helsinki.

## Legends

**Supplementary Figure 1. An outline of the study design**

*ELISA: Enzyme Linked Immunosorbent Assay, IgM: Immunoglobulin M, IgG: Immunoglobulin G, RT-PCR: Reverse Transcription Polymerase Chain Reaction NK: Natural killer cells, ADCC: antibody dependent cellular cytotoxicity*

**Supplementary Figure 2. Flow chart indicating the number of samples tested in an individual assay**

*CHIKV: chikungunya virus, ELISA: Enzyme Linked Immunosorbent Assay, IgM: Immunoglobulin M, IgG: Immunoglobulin G, RT-PCR: Reverse Transcription Polymerase Chain Reaction NK: Natural killer cells, ADCC: antibody dependent cellular cytotoxicity*

**Supplementary Figure 3. Gating strategy for antibody-dependent NK cells activation using flow cytometry**

(*A)The lymphocytes were gated using FSC/SSC scatter, (B)The NK cells were identified as CD3−CD56+ cells and assessed for surface CD107a expression and intracellular IFN-γ expression, (C) Unstimulated control, (D, E) Representative displays of ADCC non-responder and responder to CHIKV antigen, (F) Representative dot plots for positive control (purified anti-CD16)*

**Supplementary Figure 4. Association of heterologous ADCC response with anti-CHIKV IgG antibody status**

*CHIK: Chikungunya, R: responders, ADCC positive responders: Test >3*NC, Bar graphs representing % of positive heterologous ADCC responders in anti-CHIKV IgG and IgG3 antibody positive and negatives*

