## Supplementary Figures for "Natural killer cells mediated antibody dependent cellular cytotoxicity response in acute and recovery phases of chikungunya"

**Supplementary Figure 1. An outline of the study design**

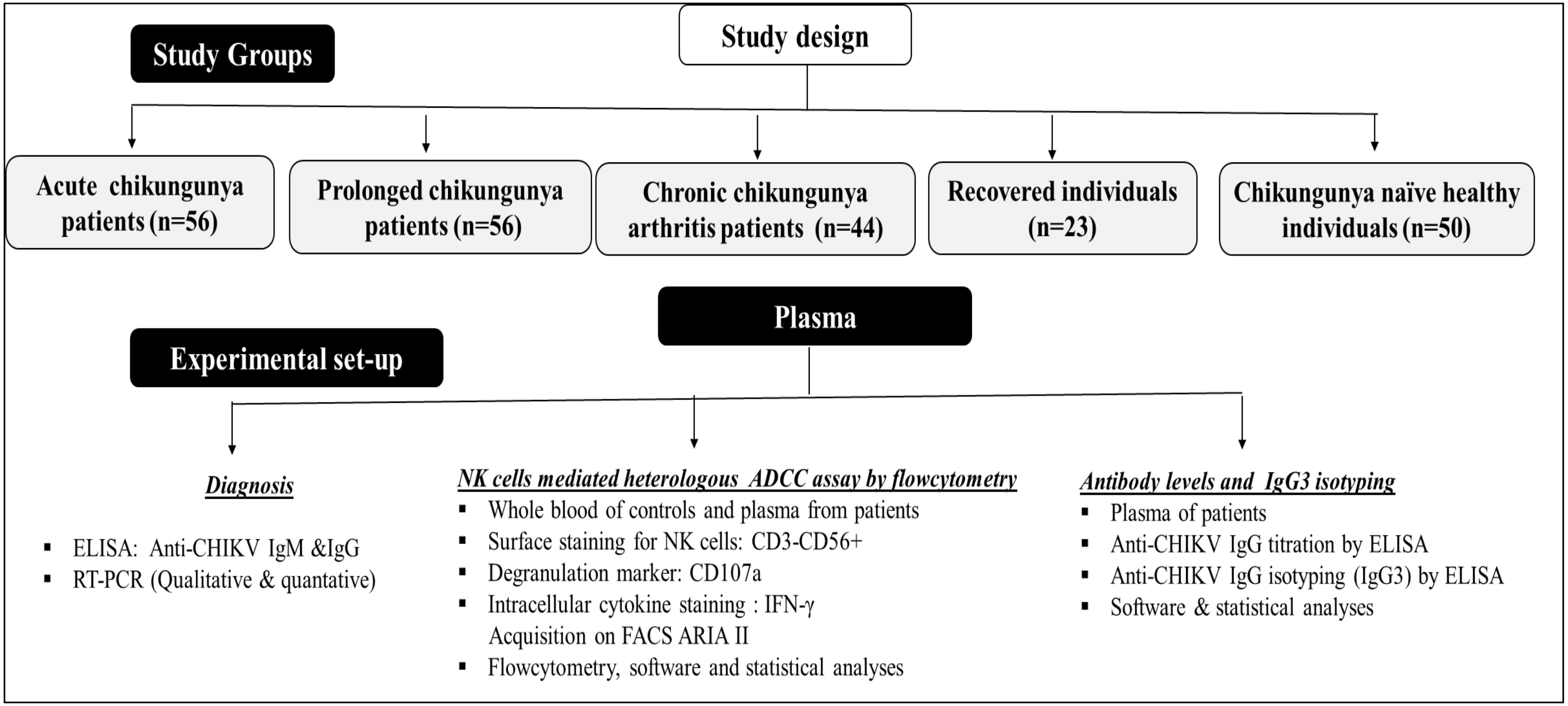


*ELISA: Enzyme Linked Immunosorbent Assay, IgM: Immunoglobulin M, IgG: Immunoglobulin G, RT-PCR: Reverse Transcription Polymerase Chain Reaction NK: Natural killer cells, ADCC: antibody dependent cellular cytotoxicity*

**Supplementary Figure 2. Flow chart indicating the number of samples tested in an individual assay**

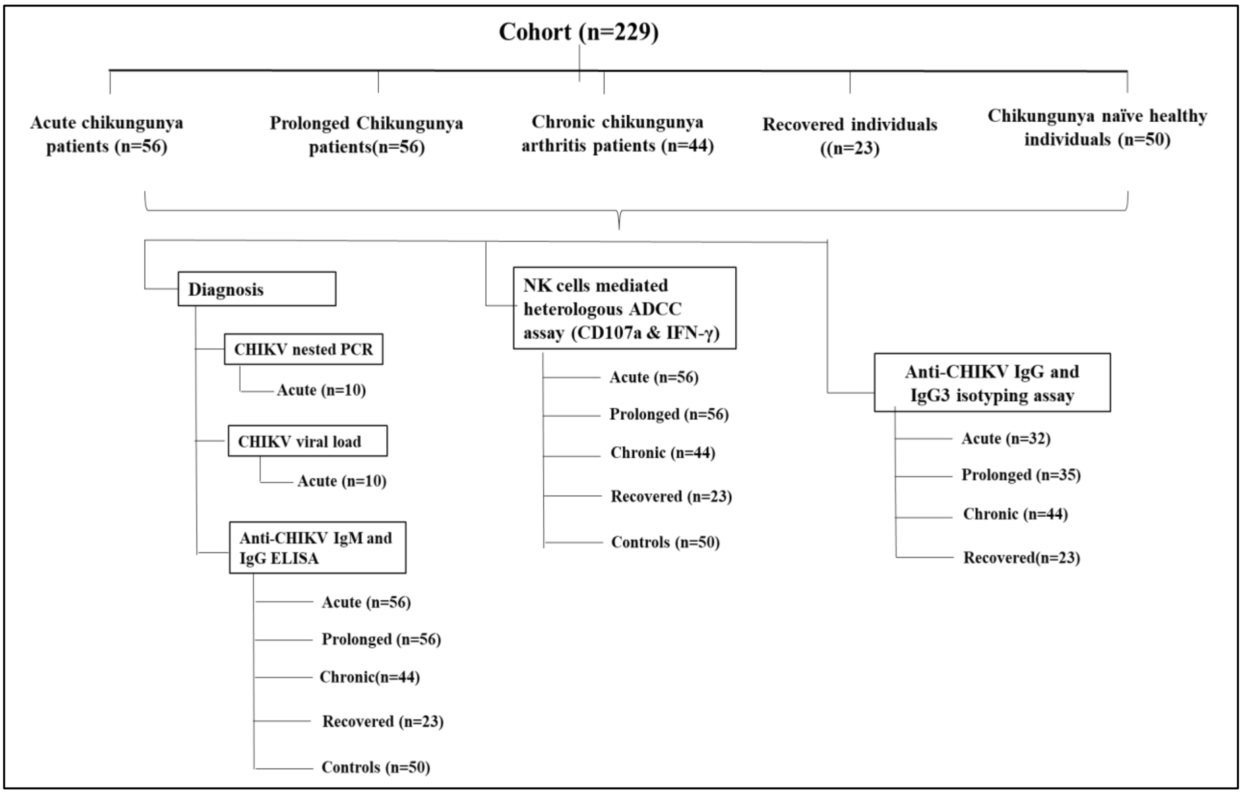


*CHIKV: chikungunya virus, ELISA: Enzyme Linked Immunosorbent Assay, IgM: Immunoglobulin M, IgG: Immunoglobulin G, RT-PCR: Reverse Transcription Polymerase Chain Reaction NK: Natural killer cells, ADCC: antibody dependent cellular cytotoxicity*

**Supplementary Figure 3. Gating strategy for antibody-dependent NK cells activation using flow cytometry**

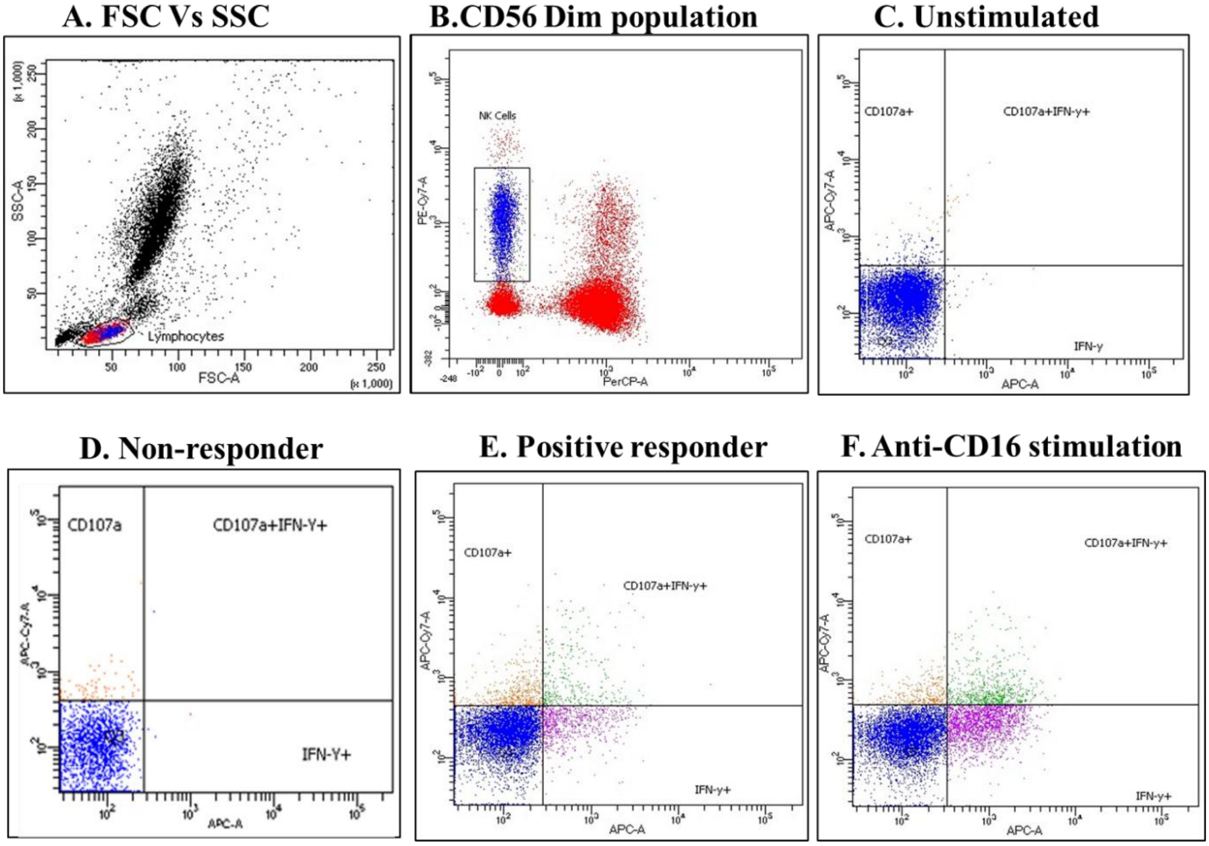


*(A)The lymphocytes were gated using FSC/SSC scatter, (B)The NK cells were identified as CD3−CD56+ cells and assessed for surface CD107a expression and intracellular IFN-γ expression, (C) Unstimulated control, (D, E) Representative displays of ADCC non-responder and responder to CHIKV antigen, (F) Representative dot plots for positive control (purified anti-CD16)*

**Supplementary Figure 4. Association of heterologous ADCC response with anti-CHIKV IgG antibody status**

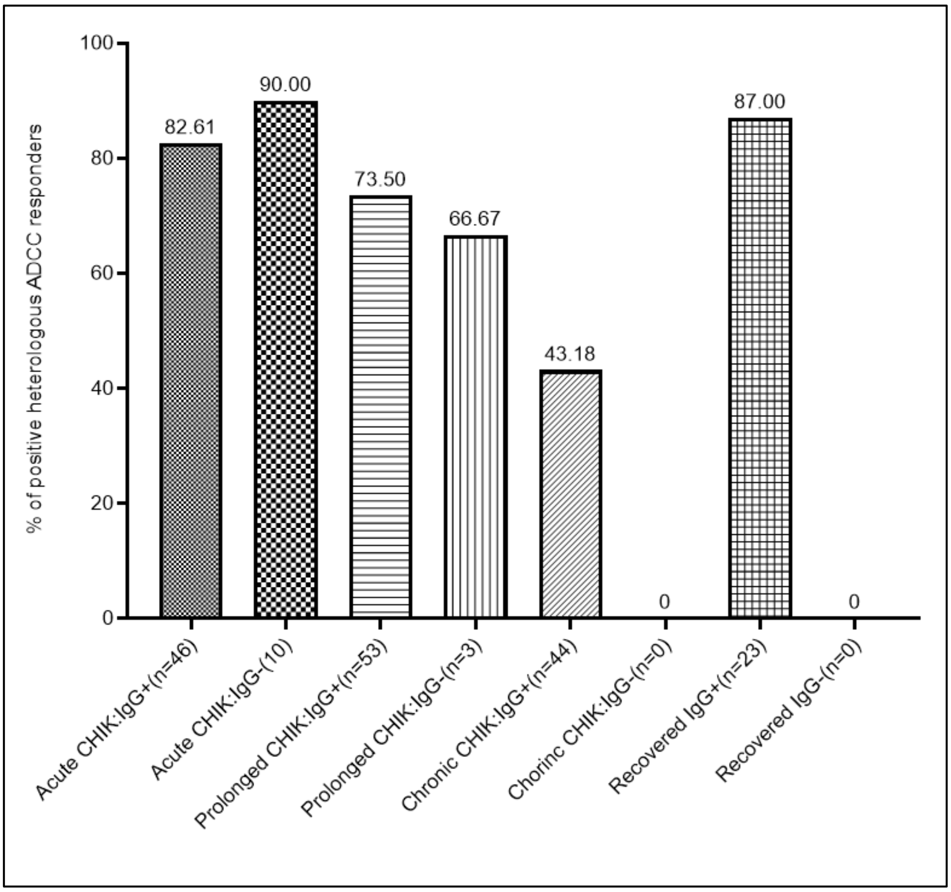


*CHIK: Chikungunya, R: responders, ADCC positive responders: Test >3*NC, Bar graphs representing % of positive heterologous ADCC responders in anti-CHIKV IgG and IgG3 antibody positive and negatives*
